# Fibre tracing in biomedical images: An objective comparison between seven algorithms

**DOI:** 10.1101/2024.04.15.589548

**Authors:** Youssef Arafat, Cristina Cuesta-Apausa, Esther Castellano, Constantino Carlos Reyes-Aldasoro

**Affiliations:** Department of Computer Science, School of Science and Technology, City, University of London, UK; Tumour-Stroma Signalling Lab, Universidad de Salamanca, Salamanca, Spain

## Abstract

Obtaining the traces and the characteristics of elongated structures is an important task in computer vision pipelines. In biomedical applications, the analysis of traces of vasculature, nerves or fibres of the extracellular matrix can help characterise processes like angiogenesis or the effect of a certain treatment. This paper presents an objective comparison of six existing methodologies (Edge detection, CT Fire, Scale Space, Twombli, U-Net and Graph Based) and one new approach called *Trace Ridges* to trace biomedical images with fibre-like structures. Trace Ridges is a fully automatic and fast algorithm that combines a series of image-processing algorithms including filtering, watershed transform and edge detection to obtain an accurate delineation of the fibre-like structures in a rapid time. To compare the algorithms, four biomedical data sets with very different characteristics were selected. Ground truth was obtained by manual delineation of the fibre-like structures. Three pre-processing filtering options were used as a first step: no filtering, Gaussian low-pass and DnCnn, a deep-learning filtering. Three distance error metrics (total, average and maximum distance from the obtained traces to the ground truth) and processing time were calculated. It was observed that no single algorithm outperformed the others in all metrics. For the total distance error, which was considered the most significative, Trace Ridges ranked first, followed by Graph Based, U-Net, Twombli, Scale Space, CT Fire and Edge Detection. In terms of speed, Trace Ridges ranked second only slightly slower than Edge Detection. Code is freely available at *github.com/youssefarafat/Trace Ridges*.

## Introduction

The morphological features of cells and their surrounding environment have been studied with interest in cancer and other conditions [1]. However, it is now recognised that the microenvironment that surrounds these cells has significant influence [2]. The extracellular matrix (ECM) plays a major role in the development of cancer [3–6] and other diseases [7]. The ECM provides essential structural support [8] to the cellular constituents of tissue. Every organ has a different ECM structure that serves its purpose. The ECM constantly changes its shape and remodels by a process known as ECM proteinases [9, 10]. Despite this, the extracellular matrix (ECM) has received less attention [11] than other factors of the microenvironment like macrophages, soluble factors or the vasculature. This could be due to the complicated 3D network of the ECM [12, 13], which is formed by many elements such as enzymes, glycoproteins and collagen.

Whilst visual observation of the extracellular matrix structure can reveal interesting characteristics, it is far too difficult to compare two populations by just observing images. Structural characteristics of the ECM such as the length, distribution, shapes and orientation of the fibre-like structure of these glycoproteins could provide valuable information about the microenvironment or treatments if these were to be objectively analysed. Quantification of the fibres and their characteristics is required and for this purpose, a robust tracing algorithm that identifies individual fibre-like structures, is necessary.

There have been many manual, semi-automatic and automatic methodologies proposed to delineate and trace elongated structures some of which are based on algorithms that can be considered more general as they can be applied in a context other than tracing, for instance, skeletonisation [14–16], watershed transform [17, 18], medial axis transform [19–21], Scale Space [22, 23] and even Edge Detection [24]. However, many of these methodologies have been proposed for tracing in specific applications, which may imply that these have been fine-tuned to the characteristics of the datasets being analysed, e.g., neuron tracing [25–27], retinal images [28–30], angiography [31–33], intravital vasculature observed with confocal, fluorescence or light microscopy [34–36], and extracellular matrix [37, 38]. In addition, deep learning architectures like the well-known U-Net [39] have been widely used in image analysis and provided excellent results without the need of hand-crafted features or algorithm. Instead, a combination of pairs of data and labels can be used to train an architecture to perform a specific task, which for the purposes of this paper is the segmentation of fibre-like structures. In this paper an objective comparison of algorithms that perform tracing of fibre-like structures is presented. Four types of datasets with very different characteristics were selected to provide a wide variety of conditions of noise and strength of the fibre structures. The images were hand-delineated to create a ground truth for objective comparisons and to provide training data for the deep learning architecture. Five existing algorithms that have been previously used for similar applications were selected: Edge detection [24], CT Fire [40], Scale Space [22, 23], Twombli [38], and Graph

Based [41, 42]. A U-Net [39] architecture was trained with pairs of patches of data and labels. In addition, a simple, yet quick and effective image processing-based tracing algorithm called *Trace Ridges* is proposed and compared with the other algorithms.

The main contributions of this work are the following:

- The objective comparison of seven fibre tracing methods. Each algorithm was tested in four images with very different characteristics. To analyse the effect of noise, three filtering approaches were applied as a pre-processing step for each of the algorithms. Three error metrics and computational processing time were extracted from each of the previous combinations.
- A new tracing pipeline called Trace Ridges was proposed. Trace ridges combines filtering, watershed transforms, edge detection and mathematical morphology to trace ridges in an image with fibre-like structures.

## 1 Materials and Methods

### 1.1 Data sets

Four biological data sets with very different characteristics were selected to compare the tracing algorithms (Fig. 1). Three data sets were selected as they had been used by the previously published algorithms and were freely available. Details of the data sets have been published previously, but a brief description for each will follow.

- Second Harmonic Generation (SHG) images of tumour bearing mouse mammary glands were published with the CT Fire algorithm [40, 43] (Fig. 1a). To obtain the images, *in vivo* acquisition was performed through a glass imaging window that was placed at proximity to palpable tumours inside the mammary glands of live 8 weeks old PyMT mice. The excitation wavelength of the SHG images was 890 nm while the emission filter was 445 nm with a 20nm bandwidth. The pulse length was roughly 100 fs. All the SHG images were verified using cellular autofluorescence from the coenzyme Flavin Adenine Dinucleotide, in slides they were verified by white light images of H&E. Images from this data set are clean and fibres are distinguishable from each other.
- Fluorescently labelled fibronectin images (Fig. 1b). Fibronectin fibroblasts were cultivated on a gelatine coated surface bonded with glutaraldehyde, that is used to create the extracellular matrix. Fluorescent images were captured to observe the fibronectin expression. Fibres appear bright and have high intensity, background is darker with lower intensity.
- Breast Cancer Biopsy (BCB) images of collagen (Fig. 1c) were published with the Twombli algorithm [38]. The samples were stained with Picrosirius red. The slides were deparaffinised and hydrated prior to the application of Picrosirius red mixture for 60 minutes, after which the samples were rinsed twice in acetic acid followed by alcohol dehydration and mounting. Finally, slides are scanned using a Zeiss Axio Scan.Z1 at 10x magnification. This dataset was chosen as the images are stained with Picrosirius red, including variety to the previous monotone images. Fibres have a filamentous structure, which make it unique.
- Disease mimicking ECM (DME) (Fig. 1d) images of fibroblasts were published with the Graph based algorithm [42]. Normal resting fibroblasts were treated with TGF-B1 cytokine to activate the fibroblast in the tumour micro-environment. The Fibronectin rich matrice was created by exposing recombinant cFn to the Fibronectin null mouse embryo fibroblasts. The cultures finally are decellularised after 7 days. The resultant matrices are viewed with a confocal microscope after they are fluorescently stained, at a 10x/0.45 magnification. Images from this data set are noisy. Brightness variation is low.

**Fig 1.**
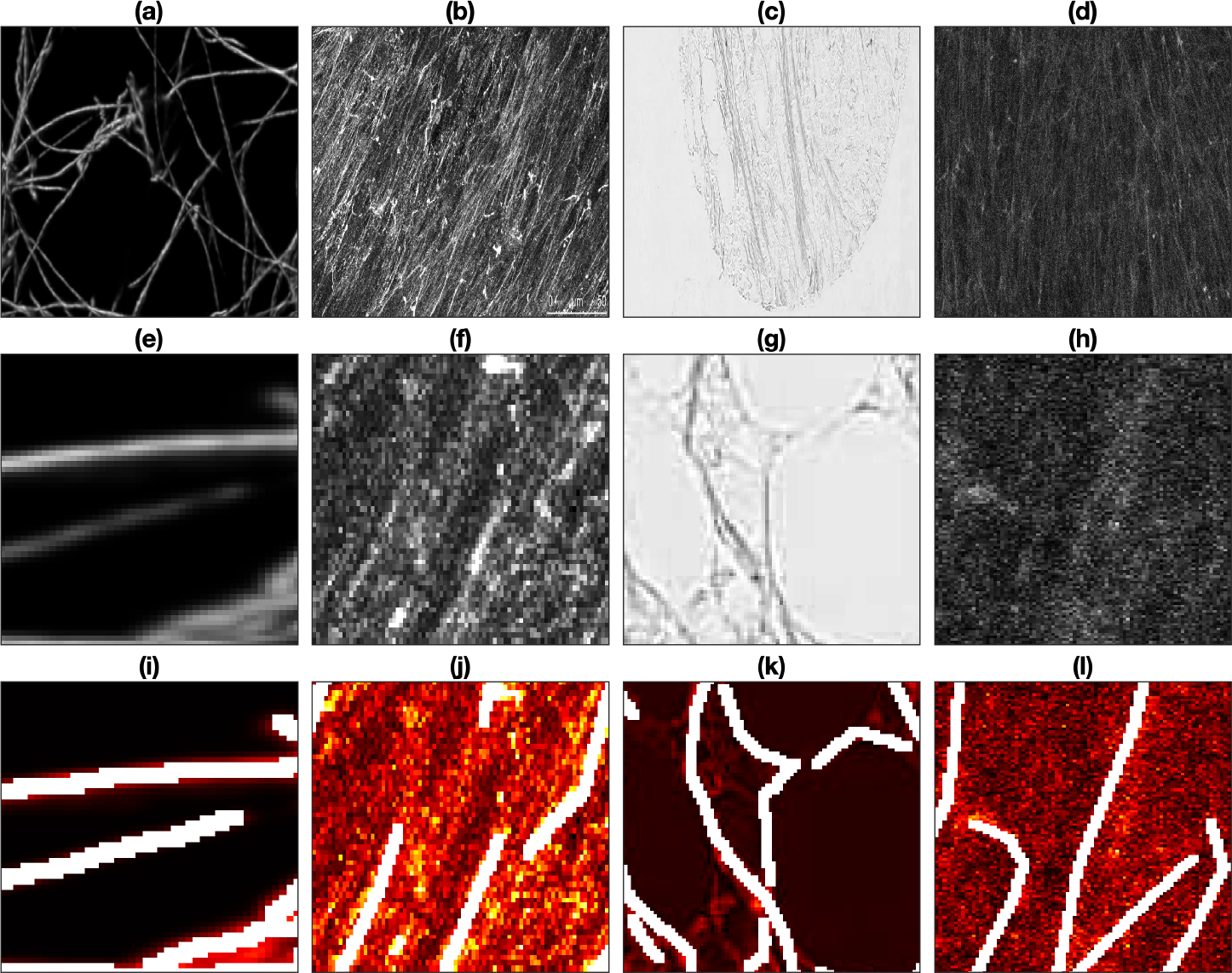
Tracing dataset(a) Second Harmonic Generation Collagen (SHG). (b) Fluorescent Fibronectin (FF). (c) Breast Cancer Biopsy slide (BCB). (d) Disease Mimicking ECM (DME). (e-h) ROI of images. (i-l) Manually delineated segmentation (GT) of ROI. White lines are the ground truth lines.

### 1.2 Pre-processing of the data sets

Filtering data sets may reduce the levels of noise and thus improve subsequent steps in a pipeline. The four data sets present very different noise characteristics, and three different filtering approaches were followed: No pre-processing filtering, low pass filtering with a Gaussian filter and a deep learning de-noising filtering [44]. It should be noted that some methodologies (e.g. Scale Space) include pre-processing steps themselves, but for consistency and ease of comparison all data sets were pre-processed in the same way before applying tracing algorithms.

#### 1.2.1 Gaussian Filtering

A Gaussian filter removes noise and detail by combining the values of neighbouring pixels weighted with a Gaussian kernel in order to replace the original value of a pixel. A Gaussian filter with a kernel size of 3 *×* 3 and a standard deviation value *σ* = 1 was applied to all the images before applying the tracing algorithms.

#### 1.2.2 DnCnn

DnCnn [44] is a de-noising deep convolution neural network for noisy images. The model adopts residual learning to separate noise from noisy observations. Batch normalisation and residual learning are adopted to improve the de-noising performance. The DnCnn model can handle blind Gaussian de-noising with unknown noise level. The DnCnn model was trained using the images in (Fig. 1). Each image was split into half, half was used for training and the other half was unseen. The images were converted to 2-D grayscale where needed as the model only accepts 2-D grayscale images. The model was trained with a learning rate of 0.001 and a batch size of 32.

### 1.3 Tracing algorithms

The following seven tracing algorithm were compared: Edge Detection (ED) [24], CT Fire (CTF) [40, 43], Scale Space (SS) [22, 23], Twombli (TW) [38], U-NET [39], Graph Based (GB) [41, 42] and the proposed Trace Ridges (TR). They vary in their complexity and methodology with some leveraging filters as part of the pre-processing (GB with Gabor filters, CTF with Curvelet transform for image enhancement). U-Net utilises deep learning techniques. Trace Ridges employs Watershed [17] and Edge Detection for fibre tracing and uses morphological features for image refinement.

#### 1.3.1 Edge Detection

The Canny Edge Detection [24] extracts edges of objects in image by identifying changes in pixel intensities. It employs Gaussian filtering to reduce noise then applies non-maximum suppression to thin the edges. Thresholding process, also known as hysteresis, is performed to exclude or include edges. In this study, the Canny Edge Detection used a filter size of 2 *×* 2 to extract edges as proxies for the fibres.

Subsequently, the edges were labelled and their metrics extracted.

#### 1.3.2 CT Fire

CT Fire [40, 43] was developed to extract and quantify collagen fibres from SHG images. It utilises a curvelet transform (CT) [45] de-noising filter, with Gaussian filtering to remove noise. The curvelet filter represents the image as stacked elements following ridge lines and wavelets, with a frequency wrapping based de-noising technique.

Curvelets commonly follow the rule in (Eq. 1). The fibre extraction algorithm processes the binary image, with foreground pixels representing fibres. It computes a distance transform to determine the proximity of the foreground pixels to the nearest background pixel, identifies maximal ridges of the smoothed image, and generates nucleation points. After which fibres extend from each nucleation point based on trajectory, followed by removal of short branches.

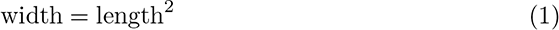

#### 1.3.3 Scale Space

Scale Space [22, 23] blurs an image across multiple scales to reveal its characteristics. Significant features emerge at lower scales, while finer details are visible at first scales and are progressively smoothed out. Scale Space treats an image as a series of Gaussian smoothed images, represented by the function f(x,y) and Gaussian g(x,y,t), where t denotes the width of the Gaussian filter. First and second derivatives of x and y dimensions are represented as

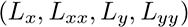

forming a matrix, dictating intensity changes in the image. For Scale Space, the collection of images is represented as L(x,y;t) and Gaussian kernel in (Eq. 2). This method was implemented as seen in [36].

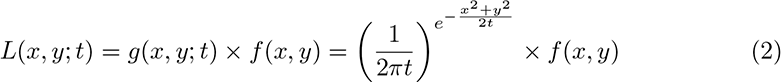

#### 1.3.4 Twombli

Twombli [38] a FIJI macro tool for quantifying fibre-like patterns. It employs existing techniques, including Ridge Detection [46] and AnaMorf [47]. Ridge Detection assesses first and second directional derivatives using Gaussian kernels to detect line points and extract line width with sub-pixel accuracy. Bias is reduced by inverting the Scale Space behaviour of the asymmetric line model. AnaMorf quantifies the structures identified. Twombli also incorporates OrientationJ to obtain the global alignment metric, determining the dominant direction of the fibrial structure. By analysing changes in intensity signal as a function of nearby pixel distances, Twombli constructs segmentation masks using fibre edges.

#### 1.3.5 U-Net

*Properties*: U-Net [39] is a deep fully convolutional network widely applied in biomedical image segmentation. The architecture compromises of a contracting path for capturing fine details through convolutions, and a symmetric expanding path for accurate localisation. The segmentation map is generated via skip connections between contraction and expanding layers. The network’s architecture is symmetrical to the letter ”U”. U-Net requires minimal training data and can achieve high performance. The U-Net used in this comparative study (Fig. 2) had a depth of 3, consisted of 46 layers, including encoder, bridge, decoder, convolution, softmax and segmentation layers, with skip connections linking the contracting and expanding paths.

**Fig 2.**
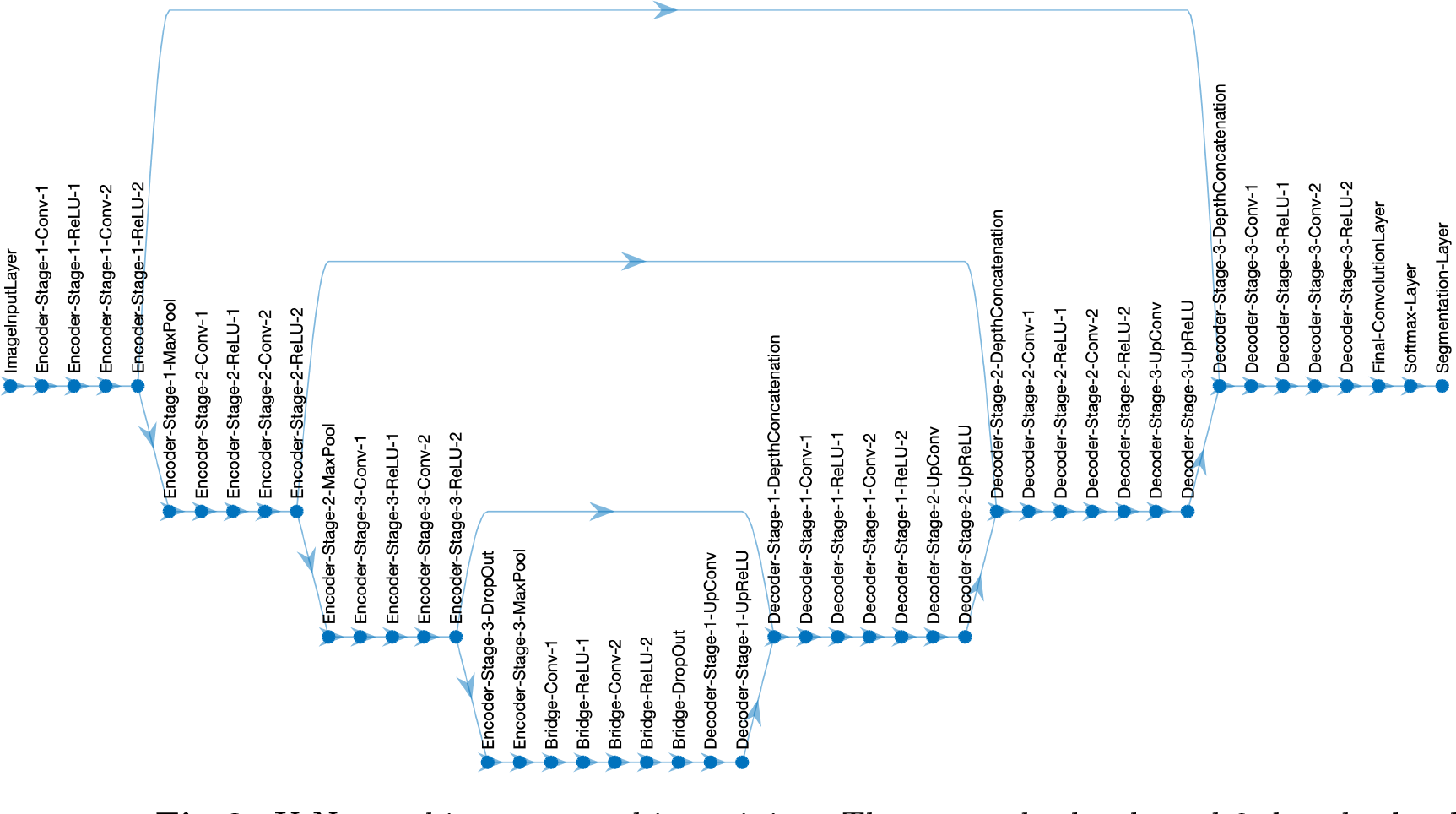
U-Net architecture used in training. Three encoder levels and 3 decoder levels. Image input layer followed by encoder, bridge, decoder, final convolution, softmax and segmentation layers.

*U-Net Training*: Initially, various training parameters were tested, including batch size (16 and 32) and learning rate (0.0001 and 0.001), to optimise training parameters and select the best-performing network. Each image from (Fig. 1a-d) was split into halves for training and testing. BCB Collagen image (Fig. 1c) was complemented to highlight fibre features in white, and background in black. To increase number of images for training, augmentation was employed. Augmentation techniques (Fig. 3) included horizontal and vertical flipping, rotation by 90 degrees and addition of Gaussian noise. A total of 8910 images of size 32x32 were used for training. The U-Net was trained with a batch size of 16, learning rate of 0.0001 and the ”adam” optimisation algorithm, for 15 epochs. Training duration was 15 minutes.

**Fig 3.**
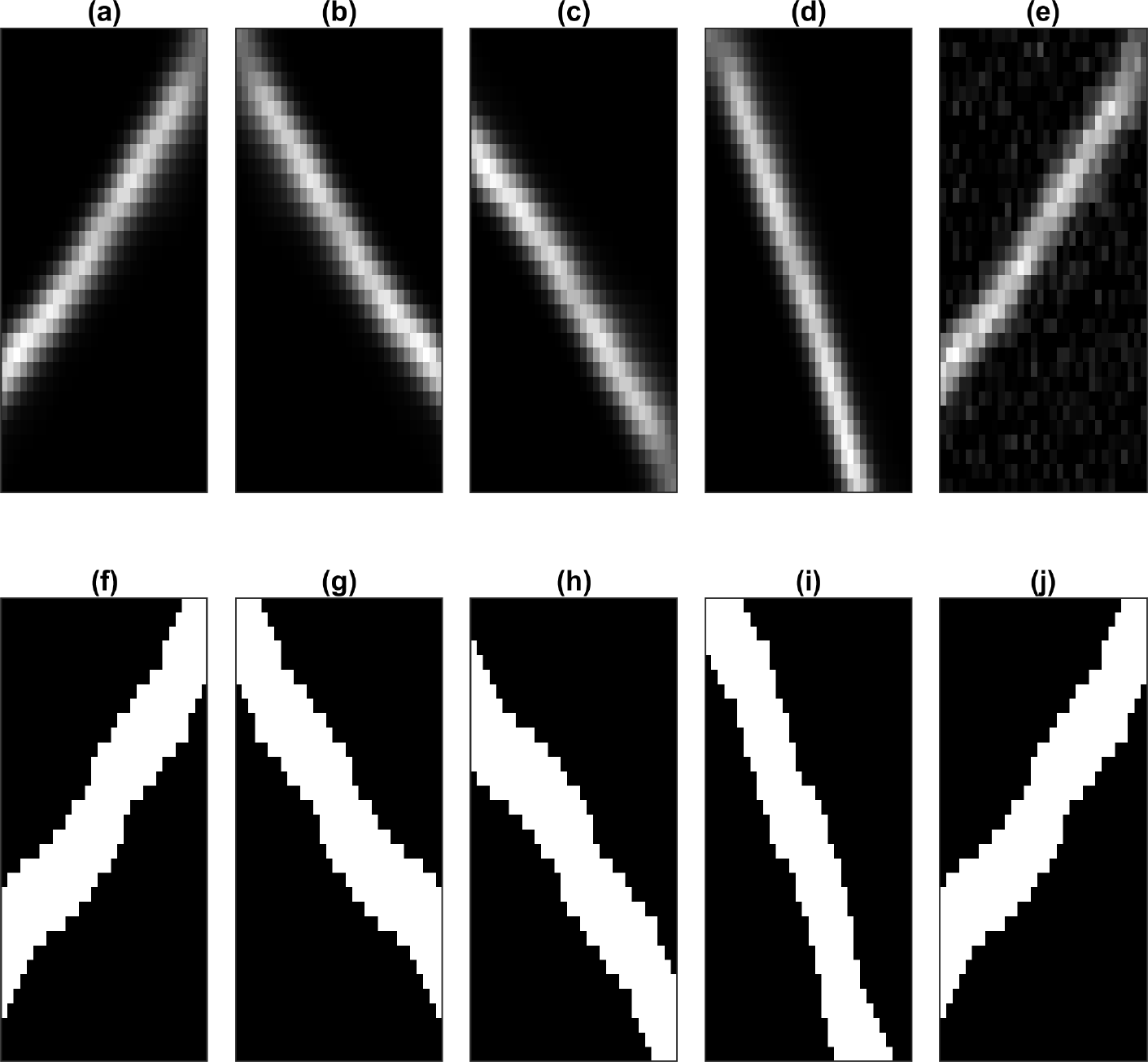
Data augmentation for training patches and their labels. (a) Original patch (b) Horizontally flipped (c) Vertically flipped (d) Rotation by 90 degrees (e) Gaussian blurred, with zero mean and a standard deviation of 0.05. (f-j) labels corresponding to augmented patches. (j) is the same as (f) as no Gaussian blur is added to the label.

#### 1.3.6 Graph Based

The Graph-based algorithm [41, 42] combines fast discrete curvelet transform filters to detect curvilinear anisotropic objects, in combination with a fibre extraction algorithm. The algorithm uses Gabor filters to enhance confocal image fibres. It offers flexibility by detecting various fibre elements at different frequencies and orientations, avoiding translation and rotation errors. The algorithm associates graph networks with fibre morphological skeletons, followed by fibre pruning and post-processing fibre re-connection. Two graph types are used to measure fibre properties: one representing fibre crosslinks or ends, and the other simplifying the first graph into a skeleton. These representations extract geometrical and topological properties from the confocal images, linking specific Gabor (example. Orientation) and graph (example. fibre length) parameters to physical fibre properties.

#### 1.3.7 Trace Ridges

In this paper, *Trace Ridges* is proposed, a method combining Watershed for ridge identification and Edge Detection which *breaks* any ridges that run from a main ridge towards the sides of the basins, separating them from the minor ones. It prioritises bright, longer ridges for cleaner results. Trace Ridges accepts images of any dimensionality, converting 3D images to grayscale and the maximum intensity projection is taken. It then identifies ridges using Watershed and edges using Canny Edge Detection with a filter size of 2 *×* 2. Edges are removed, breaking the ridges from the main ridge heading towards the sides of the basin. Distance mapping removes low-level intensity ridges while objects with a major axis length under 5 pixels are eliminated. The algorithm identifies ridges without holes based on the Euler number value of 1 and fills and thins the remaining holes. Small holes are kept while larger holes are opened. Long, straight fibres are retained, while others are broken at branches. Additionally, it checks for missed ridges ending in basins and combines identified ridges, extracting morphological metrics such as orientation, fibre count, mean intensities, and gap areas.

### 1.4 Ground truth (GT) and Quantitative Comparison

For a quantitative comparison, fibres on (Fig. 1(a-d) were manually delineated to create the ground truth.

Quantitative comparisons between the GT and results of the algorithms were calculated by first generating a distance map from the result (Fig. 4b) multiplied by the GT (Fig. 4c) and then the opposite, distance map from the GT (Fig. 4e) and multiplied against the result of an algorithm (Fig. 4f) as described in [36]. This produced two maps where the intensities of the traces corresponded to the distance from the traces of the opposite (GT/results). From those two maps the average distance error per trace, maximum distance error per trace and total distance error were calculated. Trace Ridges tracing on a SHG image (Fig. 4a) is shown in (Fig. 4d). Blue lines are that of the algorithm, red lines are from the GT, white lines are where they overlap.

**Fig 4.**
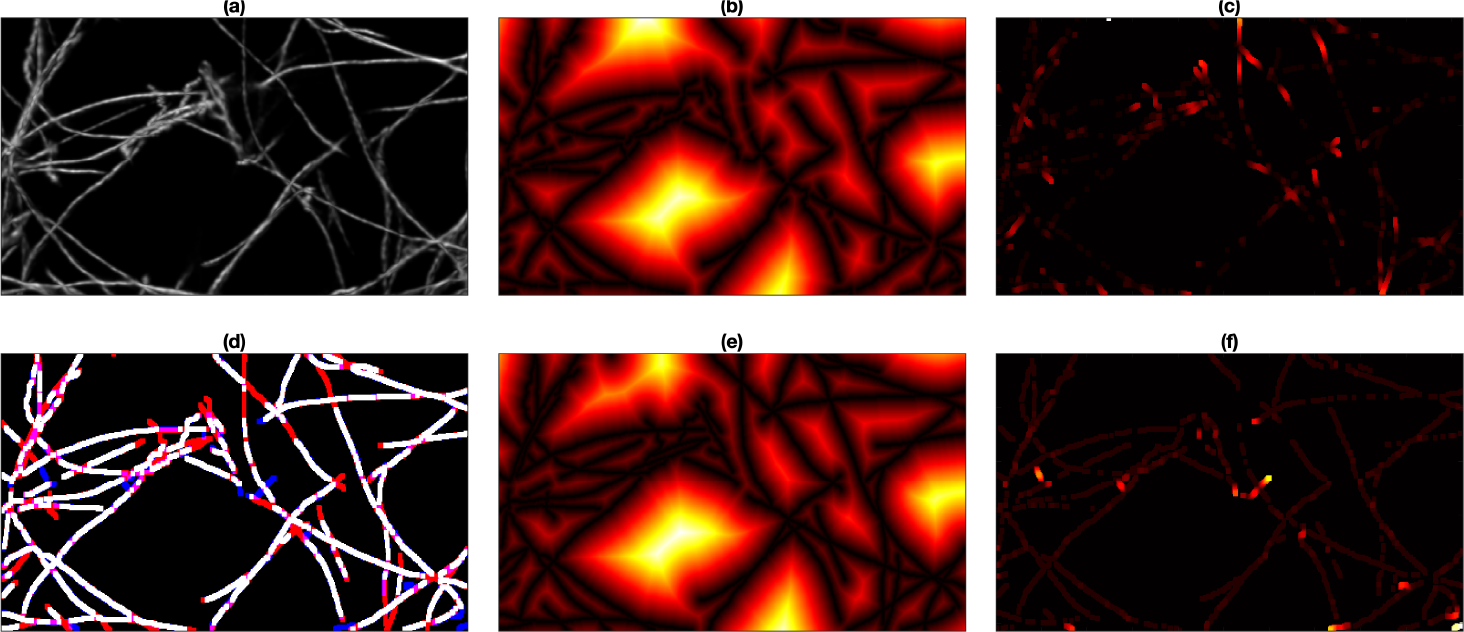
Errors distance map, pixels further away from either direction are brighter. Trace Ridges on SHG with no filtering image was used for this illustration. Trace Ridges was used as the tracing method. (a) SHG with no filtering image. (b) Distance map of the trace result. (c) GT delineation to the distance map of the trace result. (d) Tracing results of Trace Ridges. White lines indicate pixels from GT and trace. Blue lines correspond to Trace Ridges. Red lines are from the GT. (e) Distance map of the ground truth. (f) Result of the trace to the distance map of the ground truth.

## 2 Results and Discussion

In this work, images of fibre-like structures were traced by seven different image processing algorithms and were filtered by three different methods. To compare and observe which method traces fibres better, four quantitative measurements were calculated: the total distance error, average distance error and maximum distance error, as well as computational time. The distance errors are measured by comparing how close the trace results of each algorithm were compared to the manually delineated ground truth.

### 2.1 Quantitative comparison of the algorithms

#### 2.1.1 Total Distance Error

The first analysis focuses on total distance errors, this is the accumulation of the distance errors from GT to algorithm trace that are generated by the distance maps, as previously explained. Qualitative results are summarised in Table 1, comparing algorithms with and without filtering. Filtering notably improves Edge Detection, Scale Space, Twombli, Graph Based, and Trace Ridges. DnCnn enhances Edge Detection, Scale Space, and Graph Based, while Gaussian filtering benefits Twombli and Trace Ridges. However, filtering worsens Scale Space with Gaussian and Twombli with DnCnn. U-Net and CT Fire exhibit increased errors post-filtering, likely due to U-Net training on non-filtered images and CT Fire’s inherent filtering. Notably, U-Net’s performance improved when DnCnn was used on the BCB and SHG images, while U-Net with DnCnn yielded larger errors for the other images, impacting its overall performance. Considering filtering + algorithm pairs as separate methods, U-Net without filtering performs best, followed by Graph Based with DnCnn, Trace Ridges with Gaussian filtering, Trace Ridges with DnCnn and Trace Ridges with no filtering. However, averaging total distance errors across all images regardless of filtering reveals Trace Ridges as the top-performing method, followed by Graph Based, U-Net, Twombli, Scale Space, CT Fire and Edge Detection.

**Table 1.**
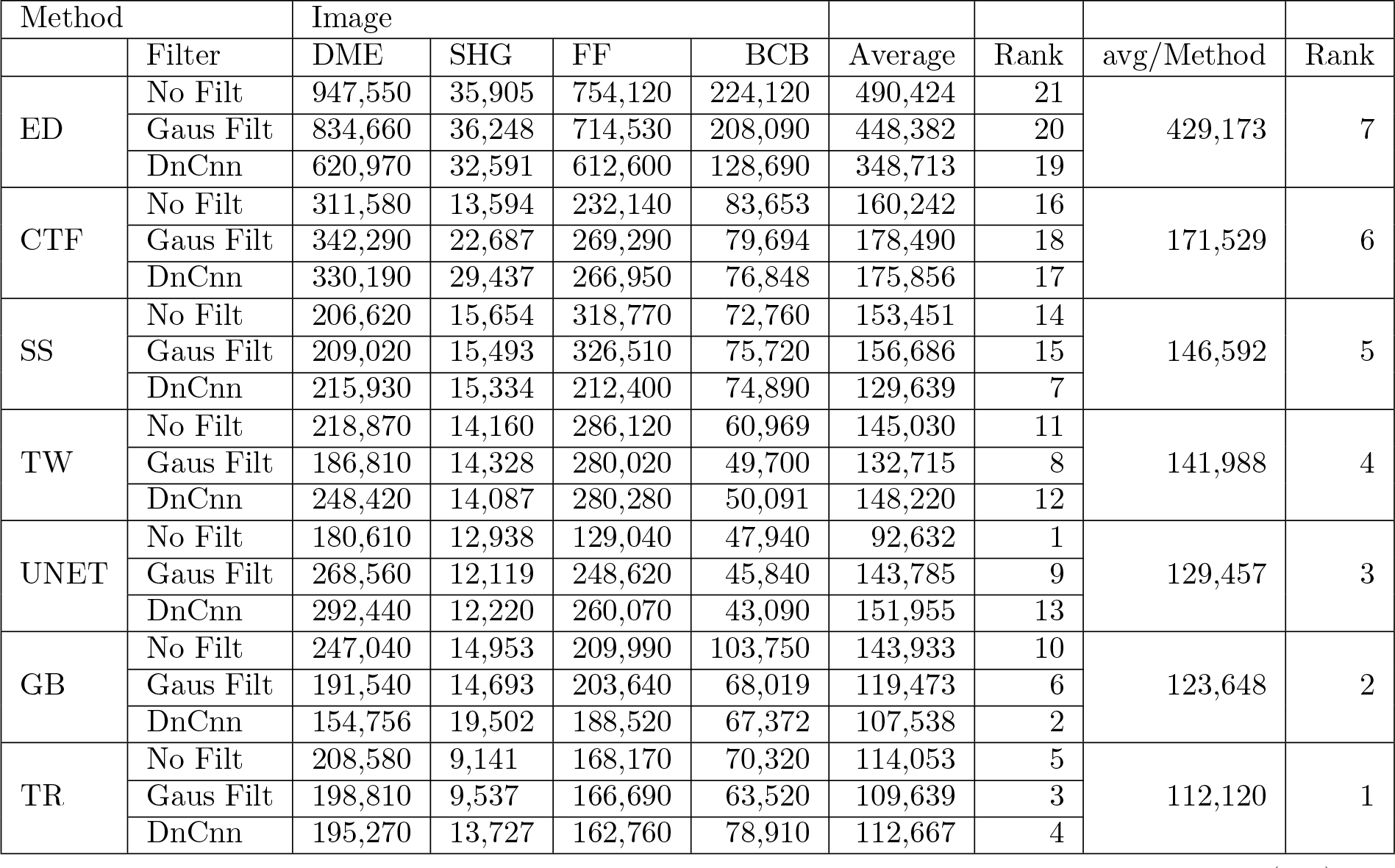
Quantitative Comparison of Tracing algorithms; Edge Detection (ED), CT Fire (CTF), Scale Space (SS), Twombli (TW), U-Net, Graph based (GB) and Trace Ridges (TR) with various filtering options on Disease Mimicking ECM (DME), Second Harmonic (SHG), Fluorescent Fibronectin (FF) and Breast Cancer Biopsy (BCB) images. Values depict total error count (pixels). Average column calculates average result for method and filter combination, followed by rank which ranks the method+filter with lower errors achieving higher ranks. avg/Method calculates average total error per method used, followed by the final rank. Algorithm result is displayed based on rank with the lowest rank at the top and highest rank at the bottom.

| Method |  | Image |  |  |  |  |  |  |  |
| --- | --- | --- | --- | --- | --- | --- | --- | --- | --- |
|  | Filter | DME | SHG | FF | BCB | Average | Rank | avg/Method | Rank |
| ED | No Filt | 947,550 | 35,905 | 754,120 | 224,120 | 490,424 | 21 | 429,173 | 7 |
|  | Gaus Filt | 834,660 | 36,248 | 714,530 | 208,090 | 448,382 | 20 |  |  |
|  | DnCnn | 620,970 | 32,591 | 612,600 | 128,690 | 348,713 | 19 |  |  |
| CTF | No Filt | 311,580 | 13,594 | 232,140 | 83,653 | 160,242 | 16 | 171,529 | 6 |
|  | Gaus Filt | 342,290 | 22,687 | 269,290 | 79,694 | 178,490 | 18 |  |  |
|  | DnCnn | 330,190 | 29,437 | 266,950 | 76,848 | 175,856 | 17 |  |  |
| SS | No Filt | 206,620 | 15,654 | 318,770 | 72,760 | 153,451 | 14 | 146,592 | 5 |
|  | Gaus Filt | 209,020 | 15,493 | 326,510 | 75,720 | 156,686 | 15 |  |  |
|  | DnCnn | 215,930 | 15,334 | 212,400 | 74,890 | 129,639 | 7 |  |  |
| TW | No Filt | 218,870 | 14,160 | 286,120 | 60,969 | 145,030 | 11 | 141,988 | 4 |
|  | Gaus Filt | 186,810 | 14,328 | 280,020 | 49,700 | 132,715 | 8 |  |  |
|  | DnCnn | 248,420 | 14,087 | 280,280 | 50,091 | 148,220 | 12 |  |  |
| UNET | No Filt | 180,610 | 12,938 | 129,040 | 47,940 | 92,632 | 1 | 129,457 | 3 |
|  | Gaus Filt | 268,560 | 12,119 | 248,620 | 45,840 | 143,785 | 9 |  |  |
|  | DnCnn | 292,440 | 12,220 | 260,070 | 43,090 | 151,955 | 13 |  |  |
| GB | No Filt | 247,040 | 14,953 | 209,990 | 103,750 | 143,933 | 10 | 123,648 | 2 |
|  | Gaus Filt | 191,540 | 14,693 | 203,640 | 68,019 | 119,473 | 6 |  |  |
|  | DnCnn | 154,756 | 19,502 | 188,520 | 67,372 | 107,538 | 2 |  |  |
| TR | No Filt | 208,580 | 9,141 | 168,170 | 70,320 | 114,053 | 5 | 112,120 | 1 |
|  | Gaus Filt | 198,810 | 9,537 | 166,690 | 63,520 | 109,639 | 3 |  |  |
|  | DnCnn | 195,270 | 13,727 | 162,760 | 78,910 | 112,667 | 4 |  |  |

**Table 2.** Quantitative Comparison of Tracing algorithms; Edge Detection (ED), CT Fire (CTF), Scale Space (SS), Twombli (TW), U-Net, Graph based (GB) and Trace Ridges (TR) with various filtering options on Disease Mimicking ECM (DME), Second Harmonic (SHG), Fluorescent Fibronectin (FF) and Breast Cancer Biopsy (BCB) images. Values depict average error distance (pixels). Average column calculates average result for method and filter combination, followed by rank which ranks the method+filter with lower errors achieving higher ranks. avg/Method calculates the average error per method used, followed by the final rank. Algorithm result is displayed based on rank with the lowest rank at the top and highest rank at the bottom.

| Method |  | Image |  |  |  |  |  |  |  |
| --- | --- | --- | --- | --- | --- | --- | --- | --- | --- |
|  | Filter | DME | SHG | FF | BCB | Average | Rank | avg/Method | Rank |
| GB | No Filt | 19.3 | 15.0 | 17.7 | 10.5 | 15.6 | 21 | 12.0 | 7 |
|  | Gaus Filt | 15.3 | 4.1 | 17.3 | 6.6 | 10.8 | 18 |  |  |
|  | DnCnn | 11.0 | 7.4 | 13.0 | 7.0 | 9.6 | 13 |  |  |
| UNET | No Filt | 10.7 | 12.9 | 10.7 | 8.2 | 10.6 | 16 | 11.0 | 6 |
|  | Gaus Filt | 11.8 | 13.6 | 14.4 | 7.5 | 11.8 | 20 |  |  |
|  | DnCnn | 12.7 | 9.7 | 14.3 | 6.1 | 10.7 | 17 |  |  |
| CTF | No Filt | 8.6 | 15.7 | 14.3 | 8.4 | 11.7 | 19 | 9.6 | 5 |
|  | Gaus Filt | 11.4 | 3.8 | 11.4 | 7.9 | 8.6 | 11 |  |  |
|  | DnCnn | 9.5 | 6.4 | 10.1 | 7.5 | 8.4 | 9 |  |  |
| SS | No Filt | 10.7 | 5.2 | 16.1 | 8.6 | 10.1 | 15 | 9.5 | 4 |
|  | Gaus Filt | 10.4 | 4.7 | 15.8 | 8.2 | 9.8 | 14 |  |  |
|  | DnCnn | 10.7 | 5.0 | 10.4 | 8.2 | 8.6 | 10 |  |  |
| TW | No Filt | 10.0 | 5.0 | 10.6 | 6.9 | 8.1 | 8 | 8.2 | 3 |
|  | Gaus Filt | 9.7 | 4.7 | 10.3 | 5.7 | 7.6 | 5 |  |  |
|  | DnCnn | 14.5 | 5.5 | 10.3 | 5.4 | 8.9 | 12 |  |  |
| TR | No Filt | 11.5 | 3.1 | 9.8 | 7.6 | 8.0 | 6 | 7.8 | 2 |
|  | Gaus Filt | 10.2 | 3.4 | 8.7 | 6.7 | 7.3 | 3 |  |  |
|  | DnCnn | 10.5 | 4.3 | 9.3 | 8.3 | 8.1 | 7 |  |  |
| ED | No Filt | 8.4 | 4.8 | 9.4 | 6.4 | 7.3 | 2 | 7.2 | 1 |
|  | Gaus Filt | 8.5 | 4.9 | 9.1 | 6.6 | 7.3 | 4 |  |  |
|  | DnCnn | 8.3 | 5.1 | 8.8 | 5.7 | 7.0 | 1 |  |  |

**Table 3.**
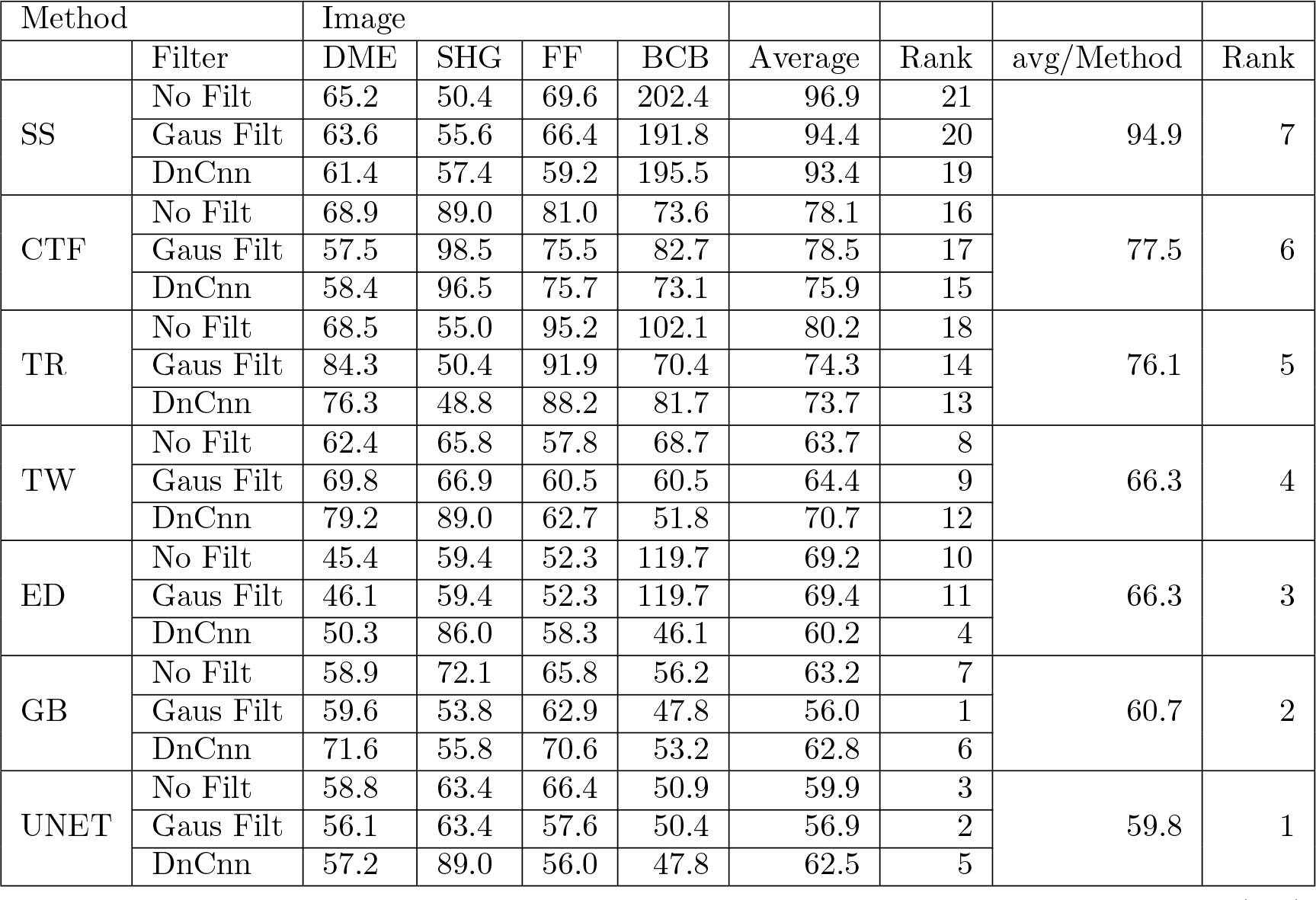
Quantitative Comparison of Tracing algorithms; Edge Detection (ED), CT Fire (CTF), Scale Space (SS), Twombli (TW), U-Net, Graph based (GB) and Trace Ridges (TR) with various filtering options on Disease Mimicking ECM (DME), Second Harmonic (SHG), Fluorescent Fibronectin (FF) and Breast Cancer Biopsy (BCB) images. Values depict maximum error distance (pixels). Average column calculates average result for method and filter combination, followed by rank which ranks the method+filter with lower errors achieving higher ranks. avg/Method calculates average maximum error per method used, followed by the final rank. Algorithm result is displayed based on rank with the lowest rank at the top and highest rank at the bottom.

**Table 4.** Quantitative Comparison of Tracing algorithms; Edge Detection (ED), CT Fire (CTF), Scale Space (SS), Twombli (TW), U-Net, Graph based (GB) and Trace Ridges (TR) with various filtering options on Disease Mimicking ECM (DME), Second Harmonic (SHG), Fluorescent Fibronectin (FF) and Breast Cancer Biopsy (BCB) images. Values depict time taken (seconds). Average column calculates average result for method and filter combination, followed by rank which ranks the method+filter with faster methods achieving higher ranks. avg/Method calculates average time taken, followed by the final rank. Algorithm result is displayed based on rank with the lowest rank at the top and highest rank at the bottom.

| Method |  | Image |  |  |  |  |  |  |  |
| --- | --- | --- | --- | --- | --- | --- | --- | --- | --- |
|  | Filter | DME | SHG | FF | BCB | Average | Rank | avg/Method | Rank |
| GB | No Filt | 1387.0 | 88.0 | 550.0 | 146.0 | 542.75 | 21 | 433.08 | 7 |
|  | Gaus Filt | 836.0 | 55.0 | 664.0 | 153.0 | 427.00 | 20 |  |  |
|  | DnCnn | 527.0 | 84.0 | 573.0 | 134.0 | 329.50 | 16 |  |  |
| TW | No Filt | 383.0 | 211.0 | 777.0 | 306.0 | 419.25 | 19 | 385.92 | 6 |
|  | Gaus Filt | 681.0 | 340.0 | 438.0 | 152.0 | 402.75 | 18 |  |  |
|  | DnCnn | 520.0 | 303.0 | 380.0 | 140.0 | 335.75 | 17 |  |  |
| CTF | No Filt | 180.0 | 60.0 | 145.0 | 77.0 | 115.50 | 15 | 102.42 | 5 |
|  | Gaus Filt | 167.0 | 57.0 | 110.0 | 71.0 | 101.25 | 14 |  |  |
|  | DnCnn | 135.0 | 49.0 | 113.0 | 65.0 | 90.50 | 13 |  |  |
| SS | No Filt | 98.6 | 4.8 | 72.7 | 45.4 | 55.40 | 11 | 55.78 | 4 |
|  | Gaus Filt | 86.0 | 5.2 | 71.8 | 82.5 | 61.36 | 12 |  |  |
|  | DnCnn | 87.0 | 5.1 | 68.0 | 42.2 | 50.57 | 10 |  |  |
| UNET | No Filt | 23.0 | 5.2 | 21.1 | 20.1 | 17.33 | 9 | 12.23 | 3 |
|  | Gaus Filt | 20.1 | 5.0 | 18.9 | 23.7 | 16.92 | 8 |  |  |
|  | DnCnn | 2.5 | 2.2 | 2.4 | 2.6 | 2.44 | 5 |  |  |
| TR | No Filt | 8.0 | 0.5 | 4.0 | 2.7 | 3.81 | 7 | 3.00 | 2 |
|  | Gaus Filt | 4.5 | 0.8 | 3.1 | 3.0 | 2.87 | 6 |  |  |
|  | DnCnn | 4.0 | 0.6 | 2.9 | 1.8 | 2.32 | 4 |  |  |
| ED | No Filt | 1.2 | 0.1 | 0.5 | 0.5 | 0.58 | 2 | 0.53 | 1 |
|  | Gaus Filt | 0.7 | 0.5 | 0.5 | 0.6 | 0.58 | 3 |  |  |
|  | DnCnn | 0.8 | 0.1 | 0.4 | 0.4 | 0.43 | 1 |  |  |

#### 2.1.2 Average Distance Error

This quantitative metric calculates total errors divided by the number of traces to provide an average, as shown in Table. 2. Filters enhance the performance of various methods including Edge Detection, CT Fire, Scale Space, Twombli, Graph based and Trace Ridges. However, filters do not improve U-Net’s values due to its training on non-filtered images. Notably DnCnn improves U-Net’s performance for the DME and BCB images, while Gaussian filtering benefits only the BCB image. Filtering yields consistent results for Edge Detection, with DnCnn slightly improving it. Both Gaussian and DnCnn filters significantly enhance CT Fire. Gaussian filtering marginally improves Scale Space, while DnCnn greatly enhances it. Twombli’s results are improved by Gaussian filtering but worsened by DnCnn, in agreement with the total distance error metric. Filtering improves the Graph based method, with DnCnn yielding the better performance. Gaussian filtering improves Trace Ridges, but DnCnn slightly increases the value, impacting performance. Filtering + method pairs are ranked, with Edge Detection + DnCnn performing best, followed by Edge Detection + no filter, Trace Ridges + Gaussian, Edge Detection + Gaussian and Twombli + Gaussian. Analysing averages per method, Edge Detection ranks highest, followed closely by Trace Ridges and Twombli. Edge Detection performs well due to the proximity of its traces to the GT, although it detects unwanted edges as fibres. Conversely, Trace Ridges, while achieving the best total distance error and the second-best average, is considered a more robust and accurate tracing method.

#### 2.1.3 Maximum Distance Error

This metric examines the maximum singular error in each image, as shown in Table. 3. Filtering enhances the performance of Edge Detection, CT Fire, Scale Space, U-Net, Graph Based and Trace Ridges. Filtering hinders Twombli. Gaussian filtering produces similar results to no filtering for Edge Detection, while DnCnn enhances results. CT Fire shows slight improvement with Gaussian and more significant improvement with DnCnn, both Gaussian and DnCnn improve Scale Space, with DnCnn yielding the lowest maximum error. Interestingly, Gaussian filtering improves U-Net but worsens with DnCnn filtering, DnCnn hinders the performance of U-Net on the SHG image, heavily impacting the average. For Graph based, DnCnn’s impact resembles that of no filtering, whereas Gaussian improves results, both Gaussian and DnCnn improve Trace Ridges, with Gaussian having a similar effect to DnCnn but slightly lesser. Among filtering + method pairs, Graph based + Gaussian performs the best, followed by U-Net + Gaussian, U-Net + no filtering, Edge Detection + DnCnn and U-Net + DnCnn. Observing averages per method, U-Net achieves the best results, followed by Graph Based and Edge Detection. However, Trace Ridges does not perform as well in this metric due to its preference for brighter and longer fibres, resulting in higher singular maximum errors. While this metric provides insights, total and average distance errors are better indicators of algorithm robustness and accuracy.

#### 2.1.4 Time

The processing time for each trace of every algorithm with various filter combinations on each image was recorded and is presented in Table. 4. Surprisingly, filtering improved the speed of every algorithm, with DnCnn consistently achieving the fastest results across all methods and filters. However, Edge Detection exhibited consistent speed with Gaussian and no filtering, and Gaussian filtering slowed down Scale Space. Each subsequent algorithm and filter pair in Table. 4 generally performed better and faster, except for Graph Based with DnCnn, which outperformed Twombli. U-Net with DnCnn was faster than Trace Ridges with Gaussian or no filtering. Edge Detection was the overall fastest algorithm, followed closely by Trace Ridges, which is known for its efficiency. U-Net was *×*3 slower than Trace Ridges, followed by Scale Space and CT Fire. Speed variations were significantly and heavy influenced by image noise levels, with noisier images (FF and DME) taking longer to process than less noisy images.

### 2.2 Visual assessment of the algorithms

A comparison between the results and the GT is presented in (Fig. 5), (Fig. 6), (Fig. 7), (Fig. 8, (Fig. 9), (Fig. 10), (Fig. 11) and (Fig. 12) with the GT in red, the results in blue. White lines correspond to pixels where the result of the algorithm and GT overlap, i.e., correct traces. It should be noted that in some cases a purple colour may be perceived, but this is an artefact when a blue line is very close to a red line.

**Fig 5.**
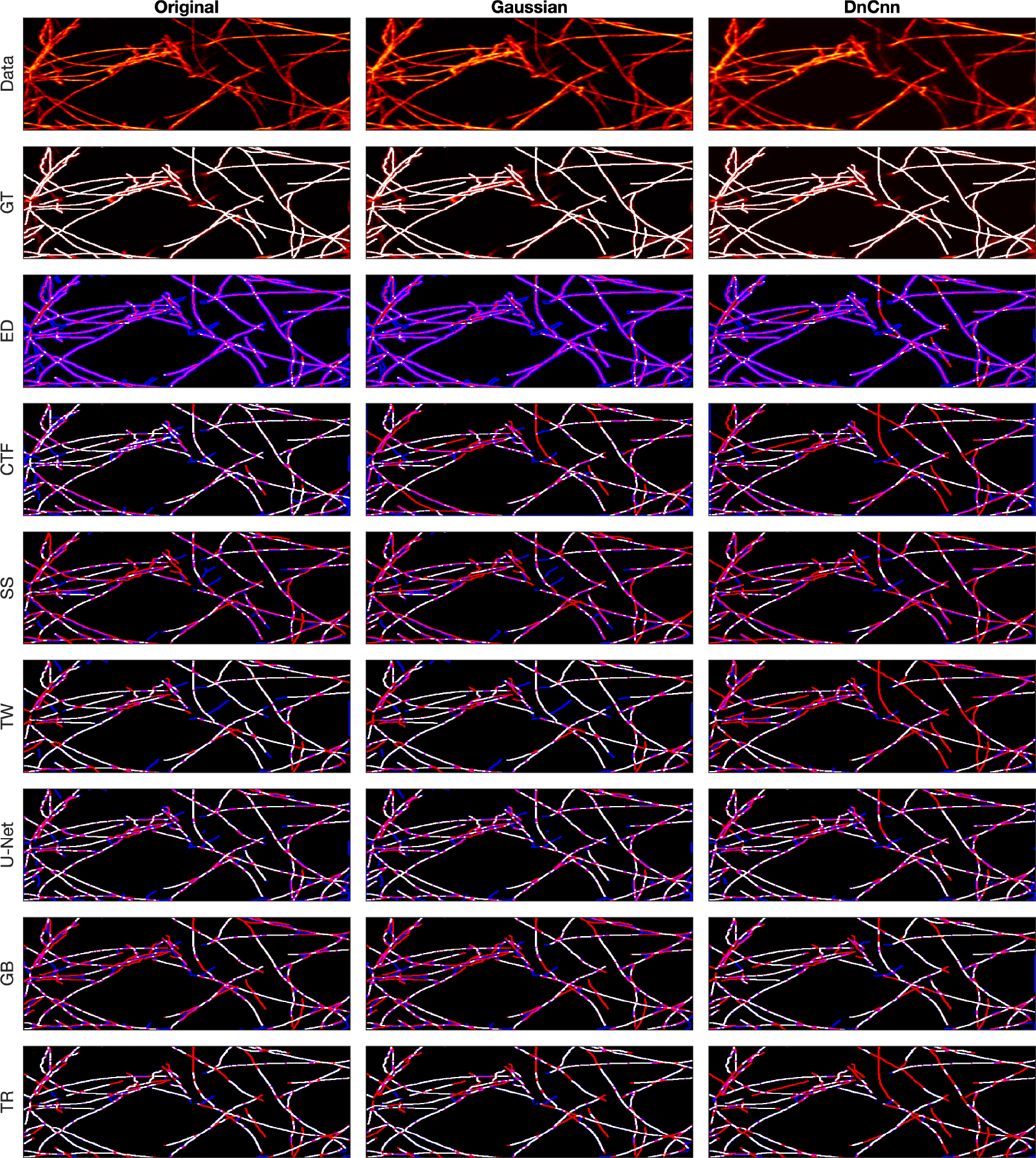
Comparison of tracing algorithms on SHG image (full view) [40]. Image was applied to algorithms with three different filters: Original/no filtering, Gaussian and DnCnn. Data corresponds to the tracing images; GT is the manually delineated ground truth. For the evaluation of Tracing Algorithms, the GT fibres are represented in red, algorithm tracing is depicted in blue and areas where they overlap is white.

**Fig 6.**
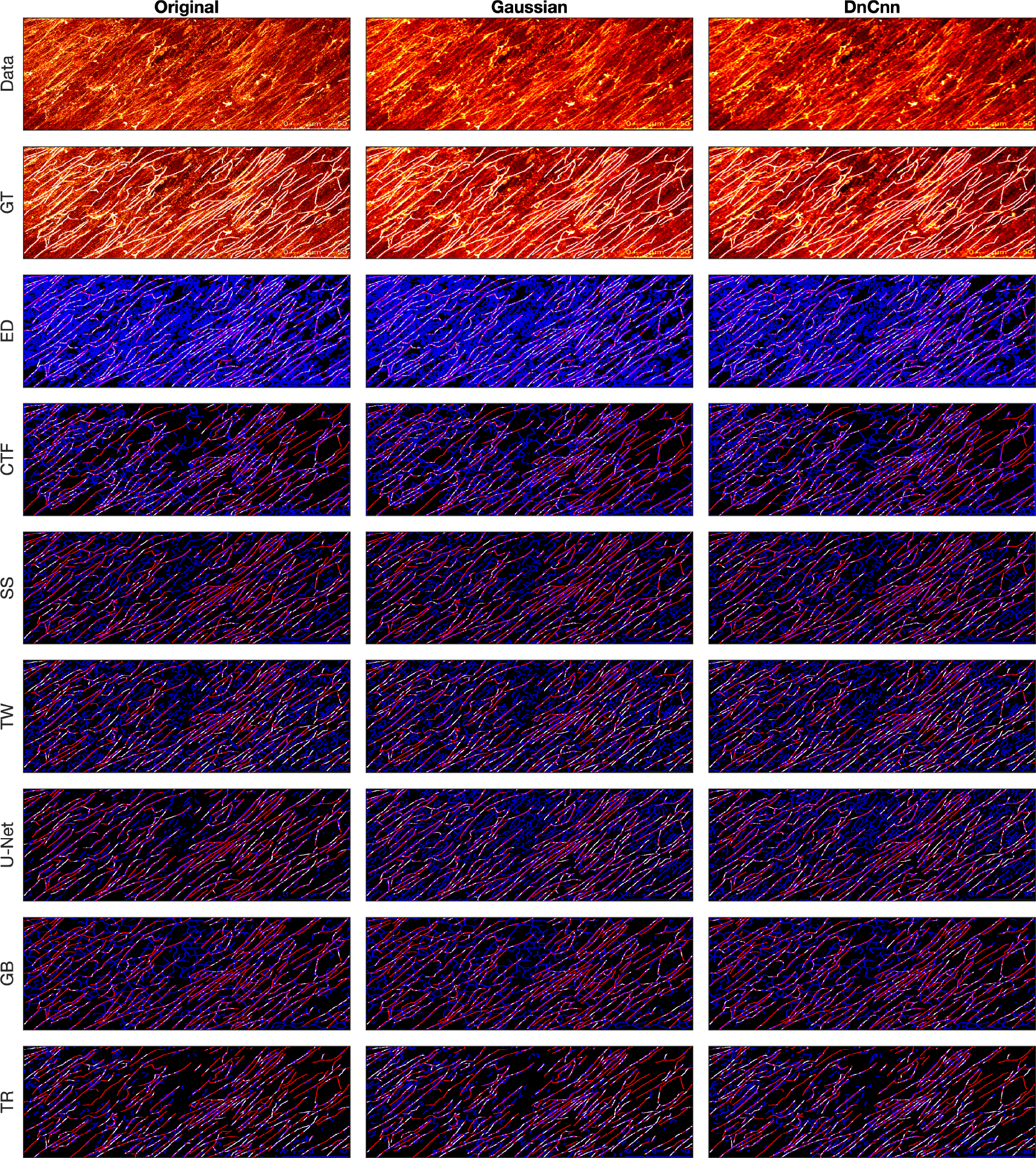
Comparison of tracing algorithms on FF image (full view). Image was applied to algorithms with three different filters: Original/no filtering, Gaussian and DnCnn. Data corresponds to the images traced; GT is the manually delineated ground truth. For the evaluation of Tracing Algorithms, the GT fibres are represented in red, algorithm tracing is depicted in blue and areas where they overlap is white.

**Fig 7.**
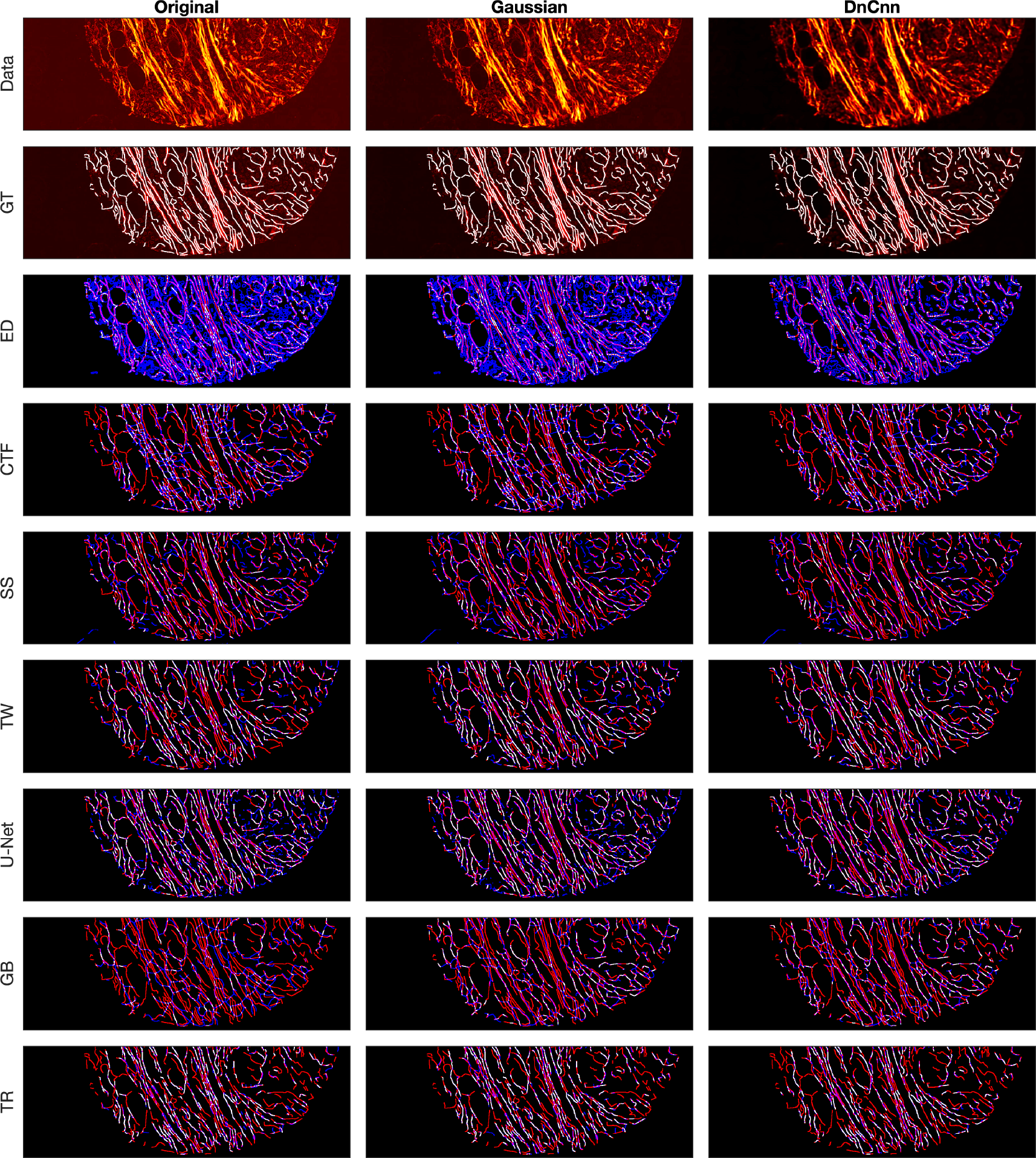
Comparison of tracing algorithms on Picrosirius red Breast Cancer Biopsy image (full view) [38]. Image was applied to algorithms with three different filters: Original/no filtering, Gaussian and DnCnn. Data corresponds to the images traced; GT is the manually delineated ground truth. For the evaluation of Tracing Algorithms, the GT fibres are represented in red, algorithm tracing is depicted in blue and areas where they overlap is white.

**Fig 8.**
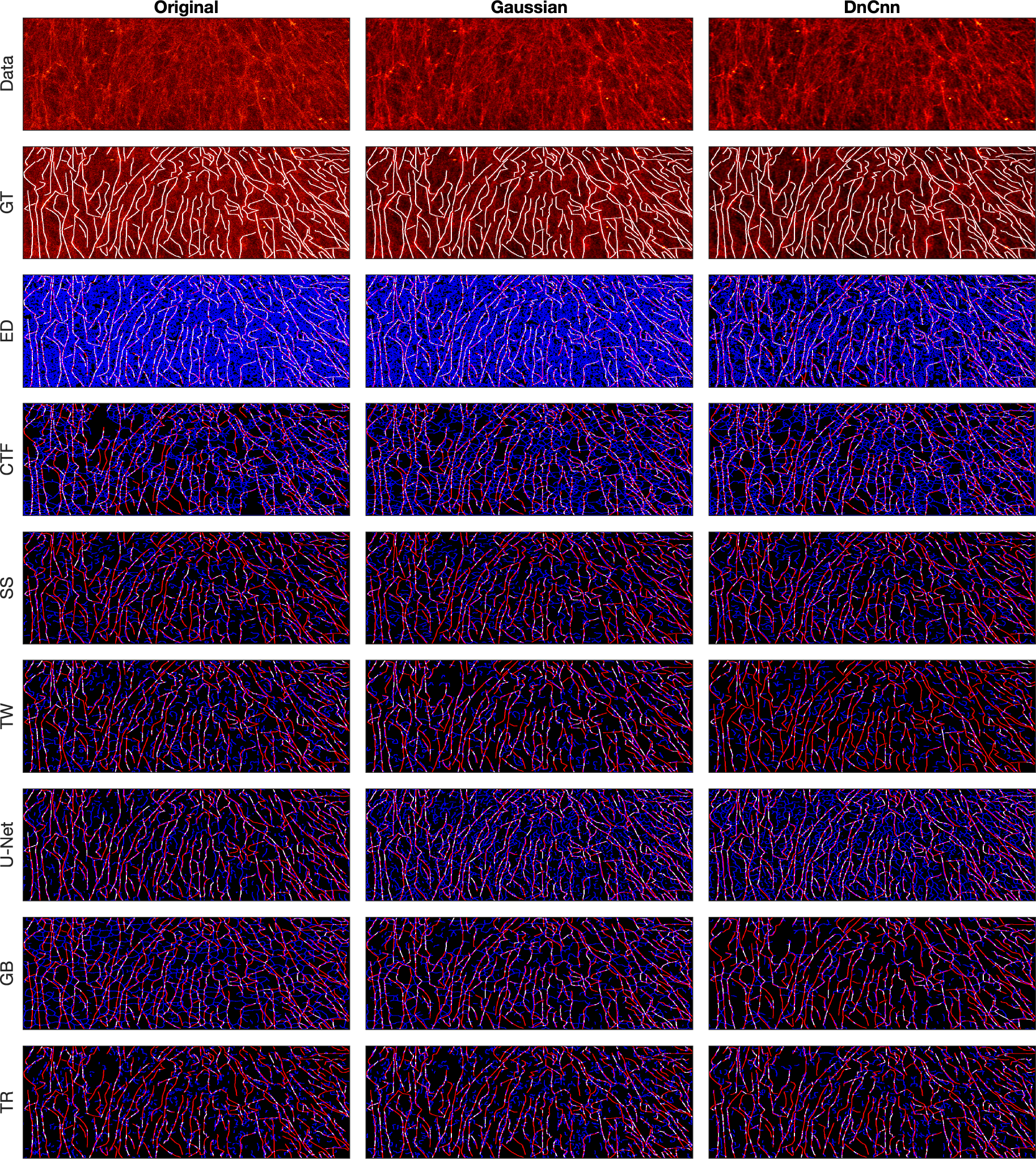
Comparison of tracing algorithms on DME image (full view) [42]. Image was applied to algorithms with three different filters: Original/no filtering, Gaussian and DnCnn. Data corresponds to images traced; GT is the manually delineated ground truth. For the evaluation of Tracing Algorithms, the GT fibres are represented in red, algorithm tracing is depicted in blue and areas where they overlap is white.

**Fig 9.**
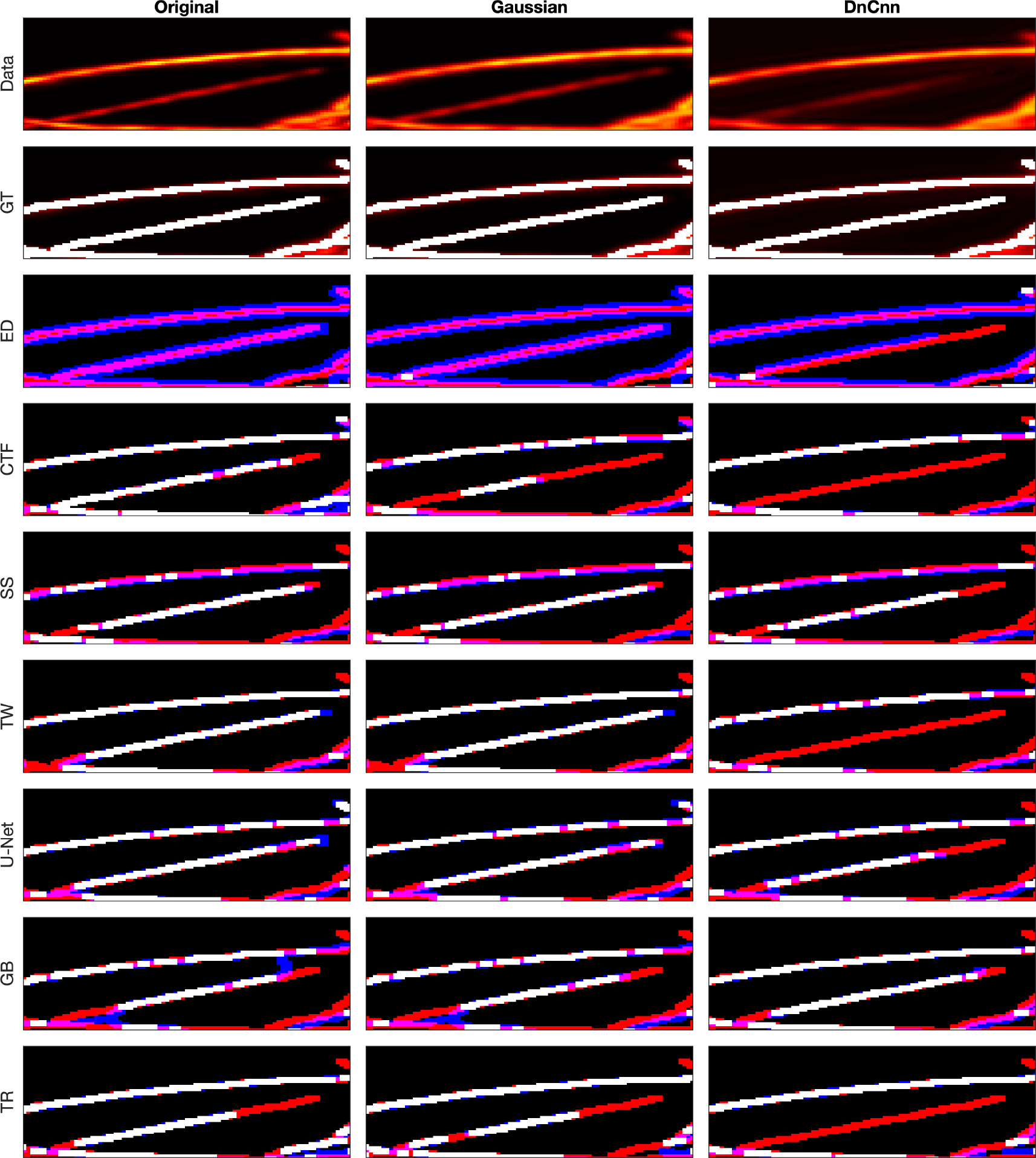
Comparison of tracing algorithms on SHG image (region of interest) [40]. Image was applied to algorithms with three different filters: Original/no filtering, Gaussian and DnCnn. Data corresponds to the images traced; GT is the manually delineated ground truth. For the evaluation of Tracing Algorithms, the GT fibres are represented in red, algorithm tracing is depicted in blue and areas where they overlap is white.

**Fig 10.**
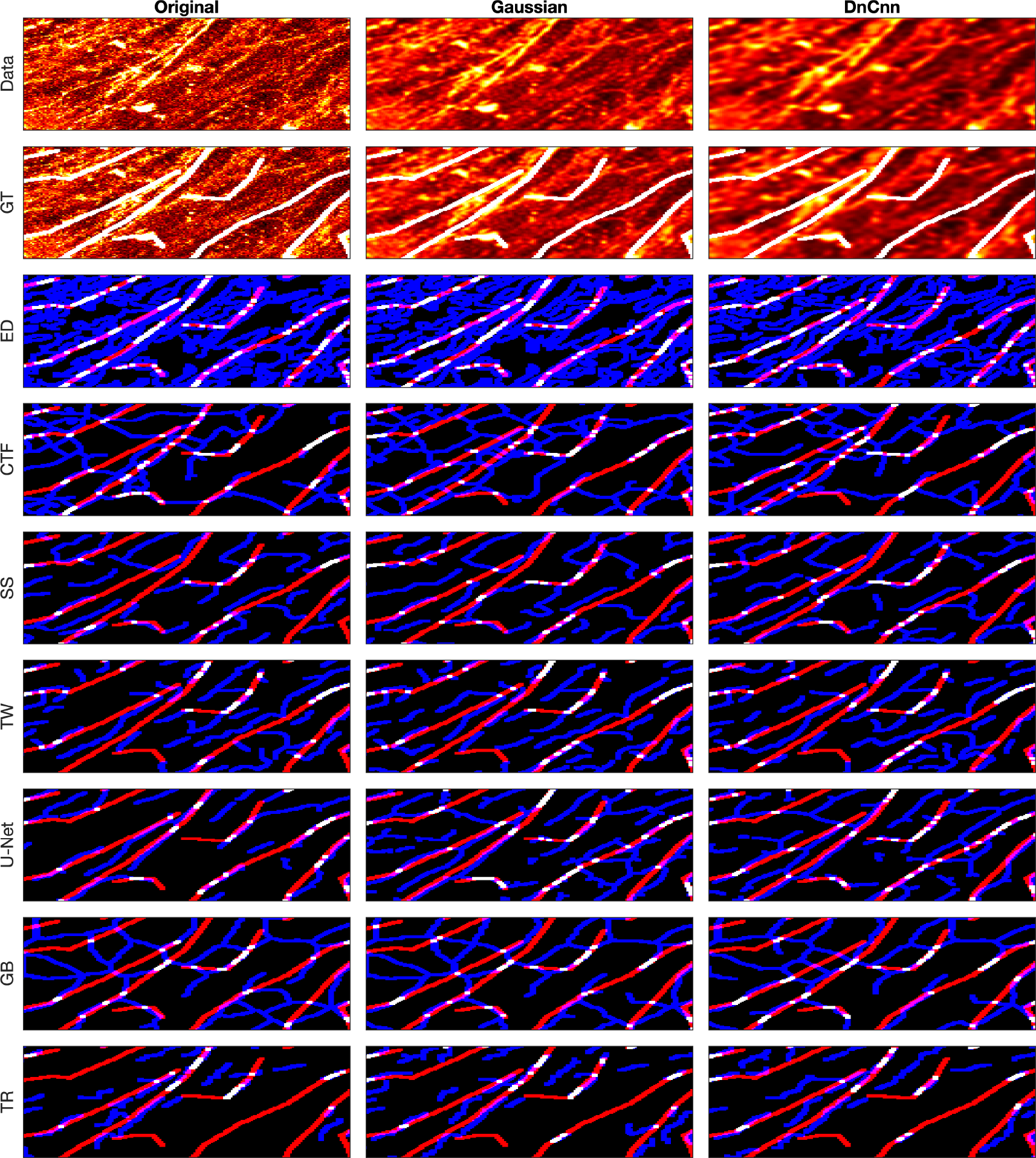
Comparison of tracing algorithms on FF image (region of interest). Image was applied to algorithms with three different filters: Original/no filtering, Gaussian and DnCnn. Data corresponds to images traced; GT is the manually delineated ground truth. For the evaluation of Tracing Algorithms, the GT fibres are represented in red, algorithm tracing is depicted in blue and areas where they overlap is white.

**Fig 11.**
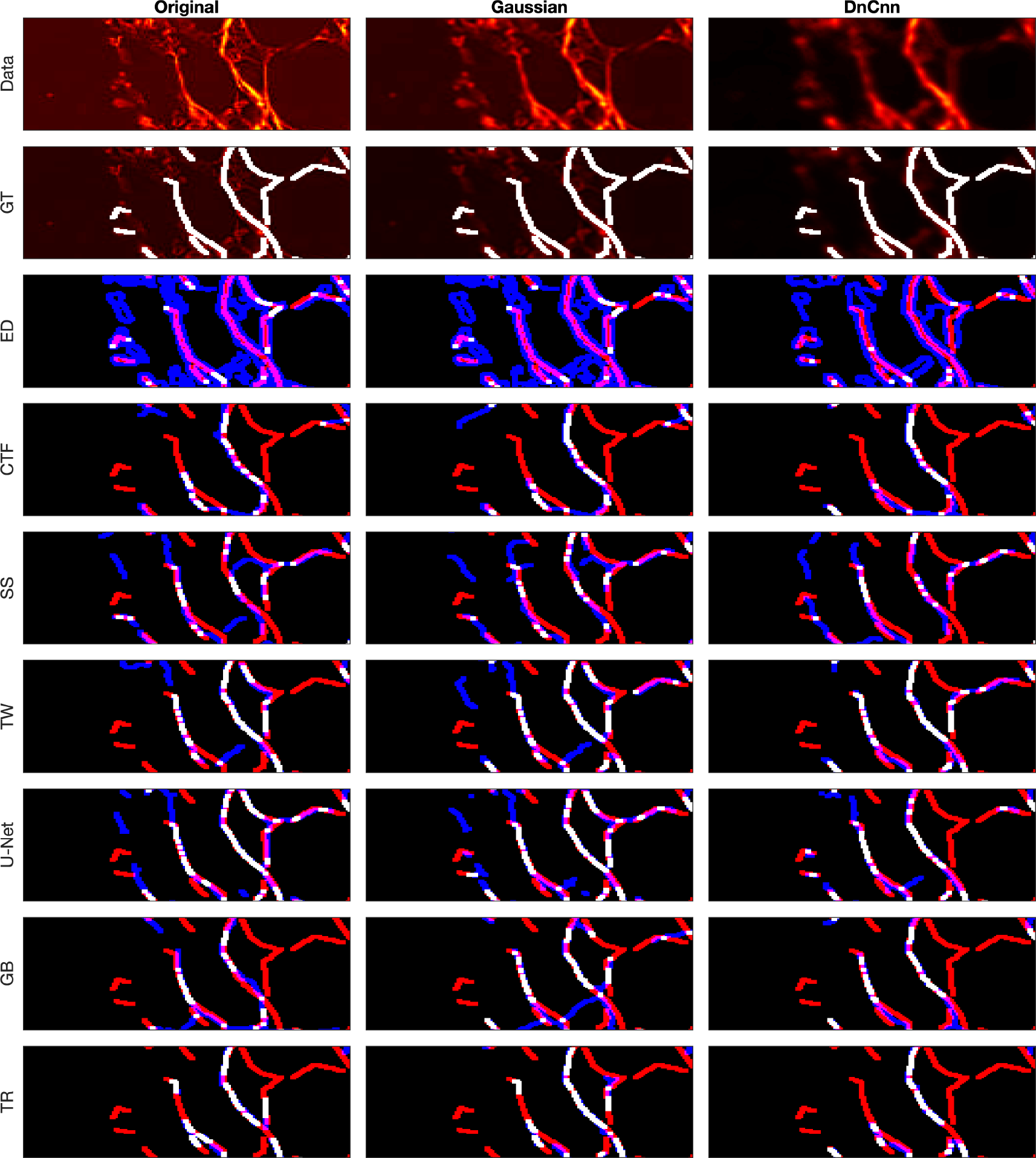
Comparison of tracing algorithms on Picrosirius red Breast Cancer Biopsy image (region of interest) [38]. Image was applied to algorithms with three different filters: Original/no filtering, Gaussian and DnCnn. Data corresponds to images traced; GT is the manually delineated ground truth. For the evaluation of Tracing Algorithms, the GT fibres are represented in red, algorithm tracing is depicted in blue and areas where they overlap is white.

**Fig 12.**
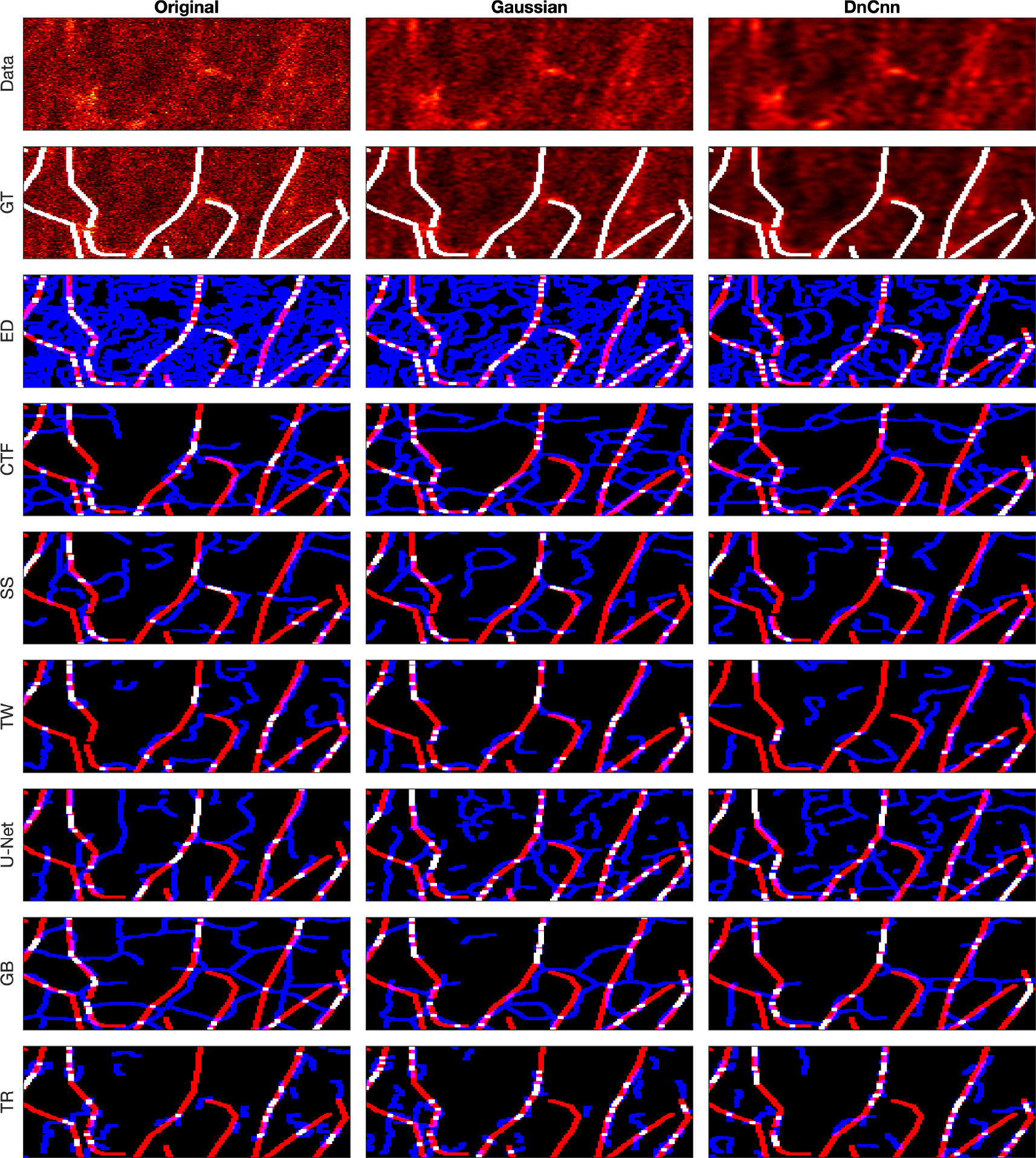
Comparison of tracing algorithms on DME image (region of interest) [42]. Image was applied to algorithms with three different filters: Original/no filtering, Gaussian and DnCnn. Data corresponds to images traced; GT is the manually delineated ground truth. For the evaluation of Tracing Algorithms, the GT fibres are represented in red, algorithm tracing is depicted in blue and areas where they overlap is white.

#### 2.2.1 Second Harmonic Generation

The SHG is a clear image, detected and missed fibres are easily identifiable. Edge Detection (Fig. 5) consistently produces duplicate lines for each fibre. From (Fig. 9), DnCnn application improves the tracing, increasing white regions compared to the original. CT Fire algorithm as demonstrated notable success on the original image (Fig. 5) but struggles with filtered images, evidenced by decreased amount of white lines regions (Fig. 9), suggesting hindrance by filtering. Scale Space remains unaffected by filtering, showing consistent results (Fig. 9). Scale Space does not perform well.

Twombli exhibits sporadic traces and misses major fibres (Fig. 5). Applicationn of DnCnn appears to worsen Twombli’s performance (Fig. 9). U-Net performs well, with significant white line regions (Fig. 5). Filtering does not visibly affect U-Net’s segmentation. Graph based method misses several fibres shown in red. It also detects spurious objects with DnCnn application (Fig. 5). Trace Ridges shows the best visual performance, with the highest number of correct fibre regions (Fig. 5). However, visually, Trace Ridges with DnCnn performs the worst among the filtering methods (Fig. 9).

#### 2.2.2 Fluorescent Fibronectin

For the FF image (Fig. 6), notable findings emerge. Despite increased noise in the image compared to the previous example (SHG), similar trends persist, with CT Fire and Edge Detection generating more traces. Filtering enhances Edge Detection (Fig. 10) but hampers CT Fire. Scale Space and Trace Ridges yield fewer traces, with Scale Space detecting darker fibres and Trace Ridges favouring the bright ones. Trace Ridges exhibits fewer spurious lines (blue) and more white regions, indicative of its strict nature (Fig. 10), showing improvement with image filtering. Graph based method and Twombli exhibit over-segmentation, visible as blue regions (Fig. 6) and (Fig. 10).

Pre-filtering notably improves their performance. Conversely, U-Net’s results with filtered images are inaccurate, as it was trained on non-filtered images, yielding optimal results with the original image (Fig. 6) and (Fig. 10).

#### 2.2.3 Breast Cancer Biopsy

For the BCB image, similar trends persist, with Edge Detection over-segmenting and producing inaccuracies, as seen in (Fig. 7) and (Fig. 11). However, pre-filtering results in cleaner images and reduces inaccuracies. CT Fire misses major fibres while detecting noise as spurious objects, evident in (Fig. 7) and (Fig. 11). Filtering enhances the algorithm’s performance, as observed in (Fig. 11). Scale Space shows inaccuracies, detecting noise and spurious objects, contrary to its usual strict behaviour, as shown in (Fig. 11). Filter application does not alter the results. Twombli shows notable performance, especially in detecting fibres at the top right corner in (Fig. 11). Gaussian filtering improves Twombli, while DnCnn hinders its performance, causing loss of white lines. U-Net’s results are intriguing, with many white regions indicating accurate delineation (Fig. 7). Filtering improves U-Net’s performance, reducing spurious objects, contrary to its behaviour on other images. Trace Ridges maintains strictness, yielding cleaner delineations (Fig. 11), albeit missing some key fibres. Graph based algorithm performs similarly with Trace Ridges, with filtering reducing spurious elements (Fig. 11.)

#### 2.2.4 Disease Mimicking ECM

A visual examination of the disease mimicking ECM (DME) image (Fig. 8) reveals notable insights. By looking at the data image, both Gaussian and DnCnn filtering visibly enhance image clarity. Edge Detection does not perform well. The main drawback of Edge Detection is that there are two lines for each fibre. The results show many blue lines, where many fibres were detected that are not there. Filtering notably enhances Edge Detection’s performance, particularly evident in the ROI (Fig. 12), where DnCnn offers the most significant improvement. CT Fire performed better than Edge Detection, it also presented spurious objects. Gaussian and DnCnn filtering resulted in more non-present traces, this is easily viewed in the ROI (Fig. 12). Scale Space displays fewer traces, benefiting from its multi resolution capabilities, but may merge separate fibres occasionally. Gaussian and DnCnn filtering have increased the number of fibres, but the latter has resulted in the long fibres to separate. Twombli yields mediocre results, segmenting non-existent fibres, yet outperforming certain algorithms. Filtering enhances Twombli’s delineation, with Gaussian filtering notably improving the algorithms performance, this is evident in the number of white lines in (Fig. 12). However, DnCnn leads to spurious lines. U-Net generally performs well, especially on the original image, but shows degraded performance on filtered images due to training on non-filtered data. The Graph based algorithm demonstrates robust performance, further improved with filtering, as seen in (Fig. 12), where DnCnn enhances accuracy and cleanliness. Trace Ridges, akin to Scale Space, exhibits fewer traces, deliberately separating intersecting fibres for improved accuracy. Filtering enhances Trace Ridges’ performance, particularly evident in (Fig. 12).

## 3 Conclusions

In this paper we introduced *Trace Ridges*, a novel method for tracing fibre-like structures, compared against six existing methods (Edge Detection, CT Fire, Scale Space, Twombli, U-Net and Graph based) with three filtering options (no filtering, Gaussian filtering and DnCnn). Gaussian filtering is a traditional filtering technique that uses Gaussian kernels while DnCnn is a deep neural network. Images representing SHG of tumor bearing mouse mammary glands images, fluorescently labelled fibronectin images, Picrosirius red breast cancer collagen fibres images and disease mimicking ECM images. These images were used for evaluation, compromising both noisy and clean samples.

Manual ground truth delineations were created for each image and distance maps were used to quantify distance errors between the ground truth and algorithmic traces.

Trace Ridges exhibited the lowest total distance error, followed by Graph based and U-Net. Edge Detection achieved the lowest average distance error, attributed to its tendency to over-segment, thus its traces are always close to a fibre from the ground truth. Edge Detection was followed by Trace Ridges and Twombli. U-Net attained the lowest maximum distance error followed by Graph based and Edge Detection. Trace Ridges did not perform well in this metric as it favours brighter and longer fibres, so it is prone to have a higher singular maximum error.

Interestingly, filtering improved results for all algorithms except U-Net, which was trained on non-filtered patches. However, filtering enhanced U-Net’s performance specifically with the Breast Cancer Biopsy image.

Edge Detection was the fastest on average, followed by Trace Ridges and U-Net.

Filtering generally expedited algorithms except for U-Net, although it improved processing time with the Breast Cancer Biopsy image.

Overall, Trace Ridges demonstrated superior performance, achieving the lowest total distance error, second-lowest average distance error, and second-fastest processing time. It offers rapid, automated tracing and extracts essential topological and morphological information, including fibre length, fibre count, orientation, circularity, and gap characteristics. Thus, Trace Ridges emerges as the preferred algorithm for tracing various fibre-like structures, including extracellular matrix and collagen fibres.

